# A unifying model that explains the origins of human inverted copy number variants

**DOI:** 10.1101/2023.09.19.558550

**Authors:** Bonita J. Brewer, Maitreya J. Dunham, M. K. Raghuraman

## Abstract

With the release of the telomere-to-telomere human genome sequence and the availability of both long-read sequencing and optical genome mapping techniques, the identification of copy number variants and other structural variants is providing new insights into human genetic disease. Different mechanisms have been proposed to account for the novel junctions in these complex architectures, including aberrant forms of DNA replication, non-allelic homologous recombination and various pathways that repair DNA breaks. Here we have focused on a set of structural variants that include an inverted segment and propose that they share a common initiating event: an inverted triplication with long, unstable palindromic junctions. The secondary rearrangement of these palindromes gives rise to the various forms of inverted structural variants. We postulate that this same mechanism (ODIRA: Origin Dependent Inverted Repeat Amplification) that creates the inverted copy number variants in inherited syndromes also generates the palindromes found in cancers.

## Introduction

Human structural variants (SVs) are DNA rearrangements that involve large segments of the human genome and can have profound effects on phenotype. They are classified as resulting in changes in copy number (copy number variants, CNVs), in chromosome location (translocations, local or dispersed duplications) and/or in orientation (direct vs. inverted). This analysis focuses specifically on locally inverted CNVs—that is, the change in copy number and orientation is confined to the vicinity of the initial region. Local inverted CNVs may include duplications and triplications, may be interspersed with copy number neutral regions, or may be associated with neighboring deletions or regions of homozygosity (ROH; also referred to as absence of heterozygosity, AOH), but have at least one repeated segment that is inverted with respect to its native orientation.

Many CNVs have been found through analysis of patients with clinical phenotypes. Identifying the genes responsible for these phenotypes is the primary focus of the clinical reports, but the sequencing data can also provide insights into the mechanism(s) responsible for their generation. The novel junction sequences that flank the CNVs have been interpreted as evidence for mechanisms of their formation and are as varied as the events themselves (reviewed in [1]). These mechanisms include breakage-fusion-bridge cycles (BFB [2]), double stranded break repair (non-homologous end joining—NHEJ [3] and microhomology mediated end joining— MMEJ [4], non-allelic homologous recombination (NAHR [5]), and altered DNA replication that result from either a collapsed replication fork (microhomology-mediated break induced replication—MMBIR [6]) or a stalled replication fork (fork-stalling and template switching— FoSTeS [7]). Sequencing of the parents’ genomes can determine whether the CNV was preexisting or *de novo*, but does not address the responsible mechanisms. Except for FoSTeS, all of these potential mechanisms have been identified, genetically dissected and verified in eukaryotic model organisms [2, 3, 5, 6]. Nevertheless, perhaps because FoSTeS is the most flexible of the models, it is frequently invoked as explaining a variety of different CNVs (for example, [8–10]). We believe that an alternative model to FoSTeS that we have experimentally validated in yeast [11–13] can cleanly explain the genesis of all locally inverted CNVs.

In this analysis, we have culled from the human literature 45 representative CNV events with localized inverted DNA segments that have well characterized junction sequences and we have explored the possibility that there is a unifying explanation for their formation. Our hypothesis is that there is a shared, unstable precursor that gives rise to the broad range of events through secondary rearrangements. That unstable structure is the palindrome: long palindromes (with perfect inverted arms), quasi-palindromes (with mismatches between the two inverted arms) and interrupted palindromes (with a short spacer between the two inverted arms) are unstable in all organisms where they have been introduced and are rare in natural genomes [14, 15] or are associated with disease [16]. Our proposal that palindromes are the predisposing structure for inverted CNVs in humans is inspired by the inverted amplification of the *SUL1* locus in the yeast *Saccharomyces cerevisiae* that invariably arises during continuous growth of laboratory strains in limiting sulfate medium. The *SUL1* amplicons have a palindromic structure with junctions that arise at closely spaced short inverted repeats. Such repeats occur at high frequency throughout the yeast genome and serve as the sites for template switching of the leading strand to the lagging strand template. Because this replication error depends on both on origin of replication and the inverted repeats, we have called this model ODIRA (Origin-Dependent Inverted-Repeat Amplification; [11–13]). Similar amplification events, consistent with an ODIRA mechanism, have also been reported at the yeast *GAP1* and *DUR3* loci [17].

While these palindromic junctions persist in yeast under selective conditions, they are unstable and can undergo secondary rearrangements through NHEJ, MMEJ, MMBIR or NAHR (Figure 1), creating new junctions and increasing the length of the spacer between the two palindromic arms. Every class of localized, inverted human CNV that we have analyzed can be explained by this same type of secondary rearrangement of an inverted triplication. Among these broad classes are inverted triplications located between direct repeats (DUP-TRP/INV-DUP), inverted duplications associated with deletions (INV-DUP-DEL), duplications/triplications flanking copy-neutral segments, absence of heterozygosity distal to the SVs (AOH), and telomere deletions distal to inverted SVs [18].

**Figure 1:**
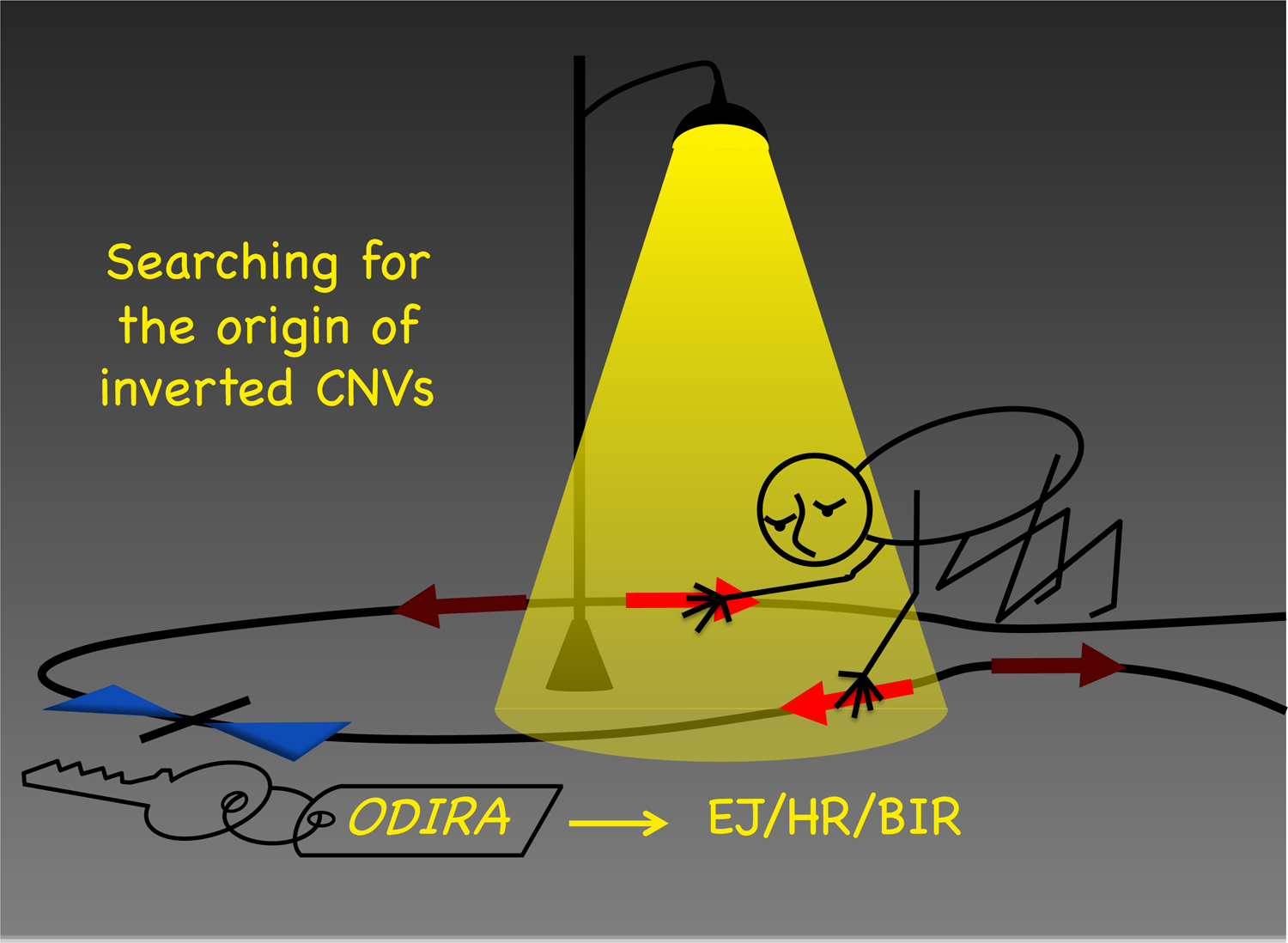
Searching for the origin of inverted CNVs. Noam Chomsky on how science operates: “Science is a bit like the joke about the drunk who is looking under a lamppost for a key that he has lost on the other side of the street, because that’s where the light is. It has no other choice.” [46] Sequencing the DNA of inverted CNVs is like looking under the lamppost. It reveals the eventual rearranged junctions (through forms of end-joining; homologous recombination or break-induced replication—EJ, HR and BIR, respectively), but may not capture the “key” initiating event (origin dependent inverted repeat amplification–ODIRA) that remains hidden in the shadows.

Genome-wide palindrome analysis (using a snap-back assay and sequencing after nuclease S1 treatment) reveals a rise in palindromic sequences in certain cancers [19–24]. “Cancer cells exhibit massive genome rearrangements, which include gene amplifications, translocations, and deletions, and these rearrangements are often associated with the presence of a palindrome, suggesting a possible correlation between the palindrome and the gene rearrangements.” In this National Cancer Institute interview Allison Rattray went on to say, “DNA palindromes are unstable and can lead to genome rearrangements by themselves, further suggesting palindromes could arise not only by sister chromatid fusion, but also by other mechanisms, such as replication errors.” (Platinum Highlight article, NCI, July 29, 2015, by Nancy Parrish, interview with Allison Rattray; https://ncifrederick.cancer.gov/about/theposter/content/novel-method-developed-further-understanding-dna-palindromes). We believe that we have identified Allison Rattray’s replication error that is responsible for many local, inverted CNVs and may be responsible for a particular subset of amplification events in cancer as well as inherited and *de novo* inverted CNVs.

## Results

Copy number variants have been routinely discovered by such techniques as array comparative genome hybridization (aCGH) or read-depth analysis of short-read sequencing data [25, 26]. However, these techniques do not reveal the genomic location of extra copies or of their orientation with respect to neighboring sequence. Discordant- or split-reads can provide information on the possible location and orientation, but are hard to definitively map in the human genome with its high density of repetitive sequences. Fluorescent *in situ* hybridization (FISH) has been invaluable for identifying and/or confirming location and orientation of duplicated or triplicated sequences; however, it does not reveal the sequences at the junctions. PCR with appropriately oriented primers can also confirm orientation at junctions. More recently, long read sequencing platforms such as PacBio and Oxford Nanopore technologies [25, 26] and Bionano optical genome mapping [27] have begun to provide much-needed tools in the identification and characterization of these local CNVs.

### Inverted Triplications

Inverted triplications with long, nearly perfect palindromic junctions (Figure 2A), such as those found at the *SUL1* locus in yeast, are difficult to identify experimentally. They can be inferred through a combination of genome wide copy number and allele frequency measurements (Figure 2B), but verification of the orientation of the amplified segments is through sequencing of the junction fragments (centromere- and telomere-proximal junctions, CJ and TJ, respectively; Figure 2). The most parsimonious structure that accommodates the number of additional copies and the junction sequences is a triplication where the center copy is inverted (TRP/INV; Figure 2A and C). Reports of TRP/INV CNVs in humans are rare, in part because older technologies using PCR and short-read sequencing are biased against recovering palindromic junctions. They are also a challenge for Nanopore sequencing [28], but as long read sequencing technologies improve, we anticipate that the frequency of inverted CNV discovery is likely to increase.

**Figure 2:**
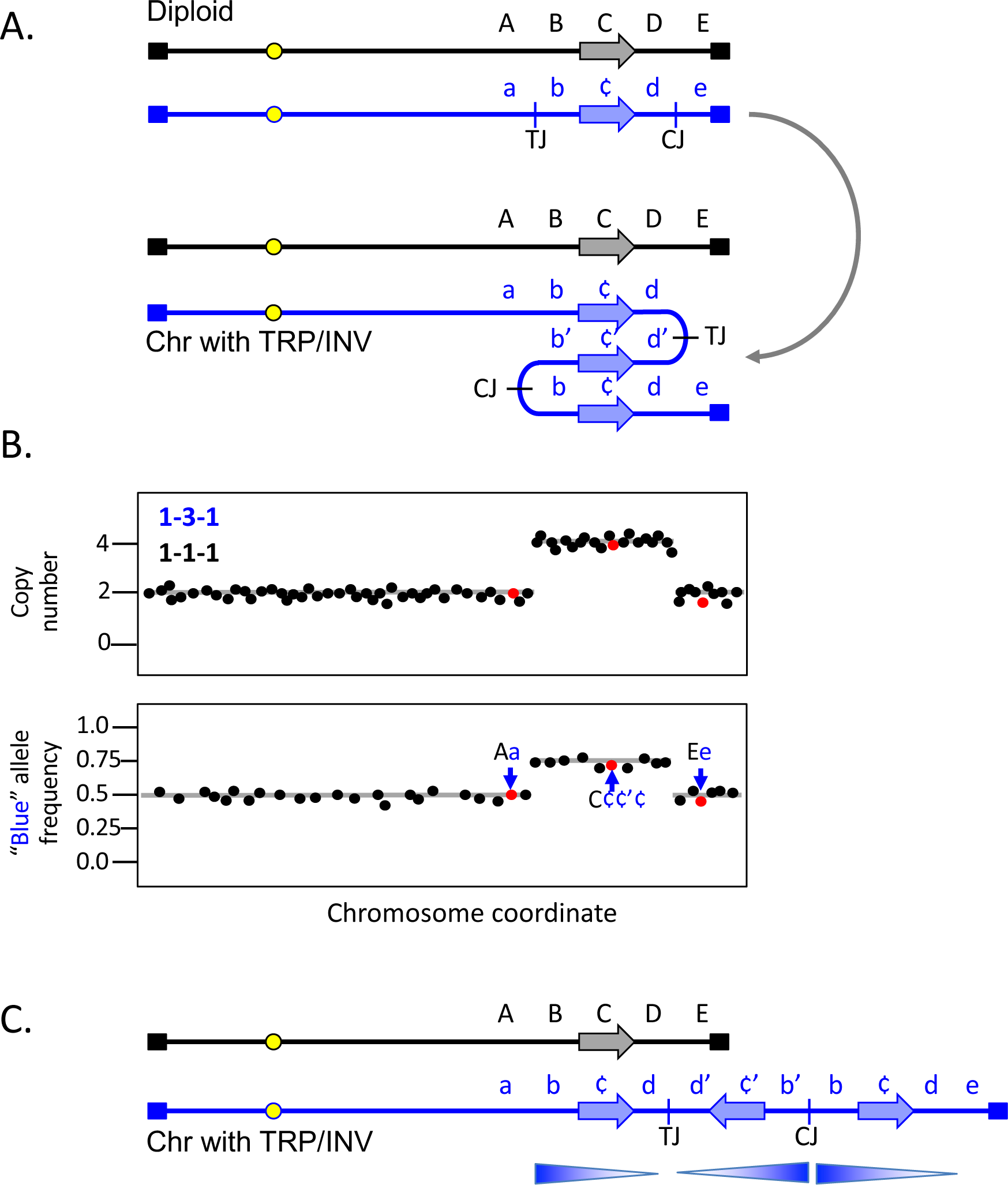
Inverted Triplication. A) A generic example of an inverted triplication in a diploid, affecting the blue chromosome, with SNPs indicated in upper and lower case letters (lower case ‘c’ being depicted as ¢ for clarity). The horizontal arrow represents a potential coding sequence. CJ and TJ refer to the potential centers of the inversion junctions (centromere-proximal and telomere-proximal junctions) identified after the inversion and triplication of the segment containing the b, c, and d SNPs. The derived chromosome is shown folded back on itself to emphasize the triplication and the inverted center copy. B) Top; expected copy number results (using either aCGH or read depth) of the diploid after triplication of the b-d region. Bottom; allele frequencies for SNPs unique to the blue chromosome. C) Linear representation of the two homologues after triplication affecting the blue chromosome. Arrows indicate the orientation of the three segments involved in the triplication. Notice that the right end of the chromosome remains intact.

Inverted triplications with their palindromic junctions are unstable [23] and secondary rearrangements that delete one of the arms of the palindromes increases the distance between inverted segments and improves their stability [29–31]. One contributing factor to palindrome instability is that repetitive elements (such as LINEs and SINEs) that were in opposite orientations before the amplification event, will have copies that are in direct orientation after the inverted duplication (Figure 3A). NAHR between two of the direct repeats would delete the segment between them (Figure 3A). This recombination event results in loss of one of the copies of DNA between the two repetitive elements and is marked by a decrease from 4 to 3 in copy number measurements and alters the allele frequencies (Figure 3B). If a similar event occurs at the other junction, then the triplication is flanked by duplications on both margins.

**Figure 3:**
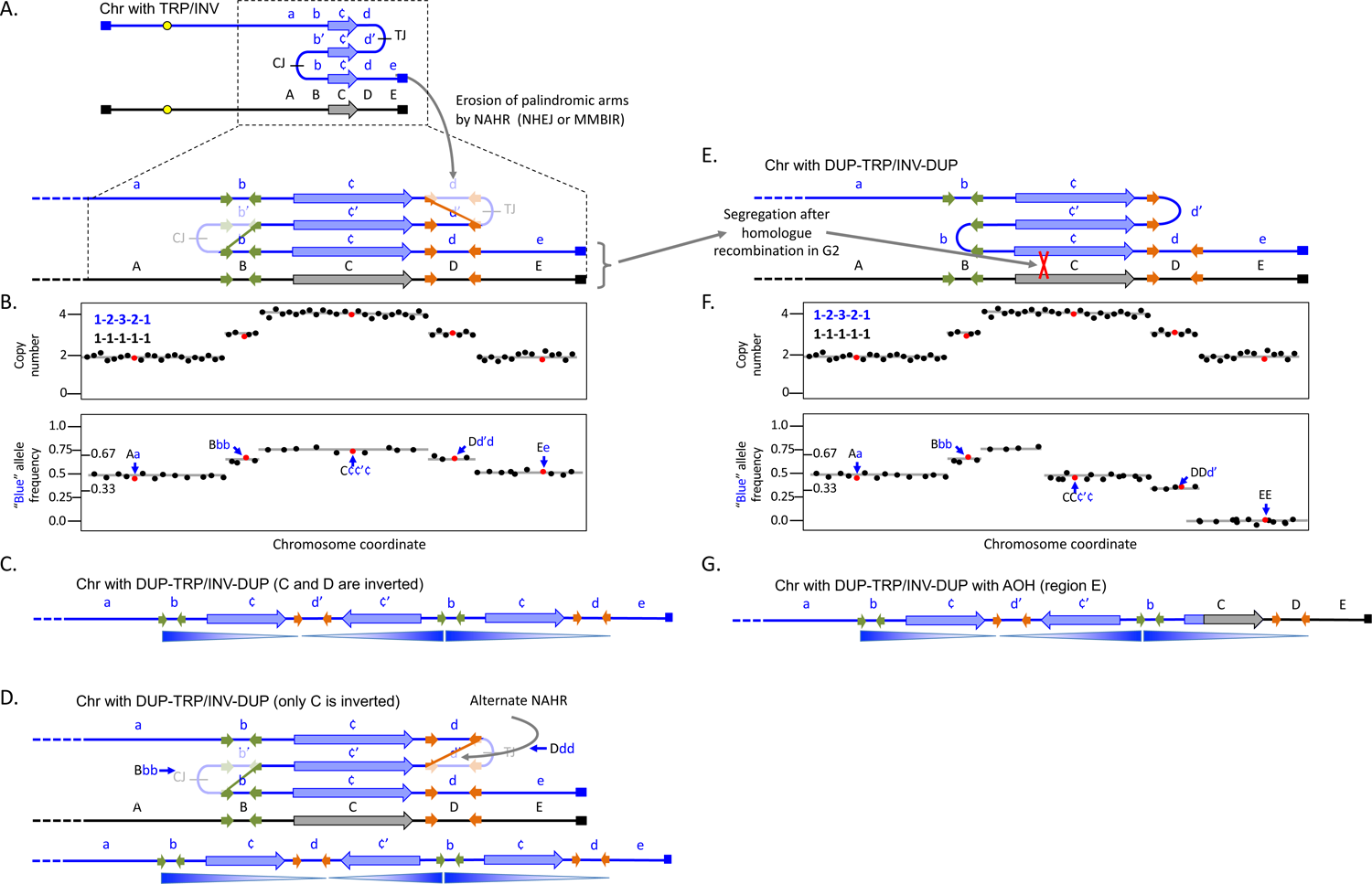
DUP-TRP/INV-DUP with and without adjacent AOH. A) The same chromosome illustrated in Figure 2 is expanded to show potential short regions of inverted homology such as SINES or Alu sequences (green and orange horizontal arrows). After the triplication, pairs of orange and green repeats are now found in direct orientation and serve as the sites for non-allelic homologous recombination or other forms of rearrangement. The original inverted junctions (CJ and TJ) have been lost and the region between the recombined repeats is reduced in copy number. B) Top; expected copy number results (using either aCGH or read depth). Bottom; allele frequencies for SNPs unique to the blue chromosome. Notice that the d region is now at three copies total with one copy of the c and d regions in inverted orientation. C) Linear representation of the blue homologue after triplication. D) An alternate recombination event at the orange repeats produces a chromosome with the same copy number and allele frequency profiles, but in this case the only the c region is inverted. E and F) After the erosion of palindromes shown in (A), a secondary event of mitotic homologous recombination and subsequent segregation produces the same pattern of copy number estimates, but the telomeric region has become homozygous for the black chromosome E allele and other c and d SNPs have been reduced. G) Linear representation of the blue homologue after homologous mitotic recombination that replaced the end of the rearranged chromosome with alleles from the black homologue. See Figure S1 for alternate illustrations of the rearrangement events shown in A and D.

For clarity, we have illustrated NAHR occurring at both junctions, but most interstitial triplications with this DUP-TRP/INV-DUP structure (Figure 3C; Table 1 examples 1-21) do not have junctions that map to LINEs, SINEs or other low copy repeats (LCRs) and are likely created through NHEJ, MMEJ or MMBIR at sites of little to no microhomology. Some notable examples of recurrent inverted CNVs have one junction between closely spaced repeats (with the alternate allele composition illustrated in Figure 3C or in Figure 3D) and the other junction created through microhomology or blunt ended ligation [32, 33]. The sizes of the DUP segments in the cases we examined ranged from as little as a few kb to thousands of kb. The triplicated segments also had a wide range of sizes. We found no examples in our limited survey where the presumptive palindromic arms were unambiguously intact. This finding is in contrast to what we recover for the yeast *SUL1* locus where only 3 of 92 sequenced junctions were consistent with secondary rearrangements [13], possibly reflecting differences in the number of cell divisions, selective pressures and/or availability of different DNA repair pathways in the yeast experiments compared to human development.

**Table 1:**
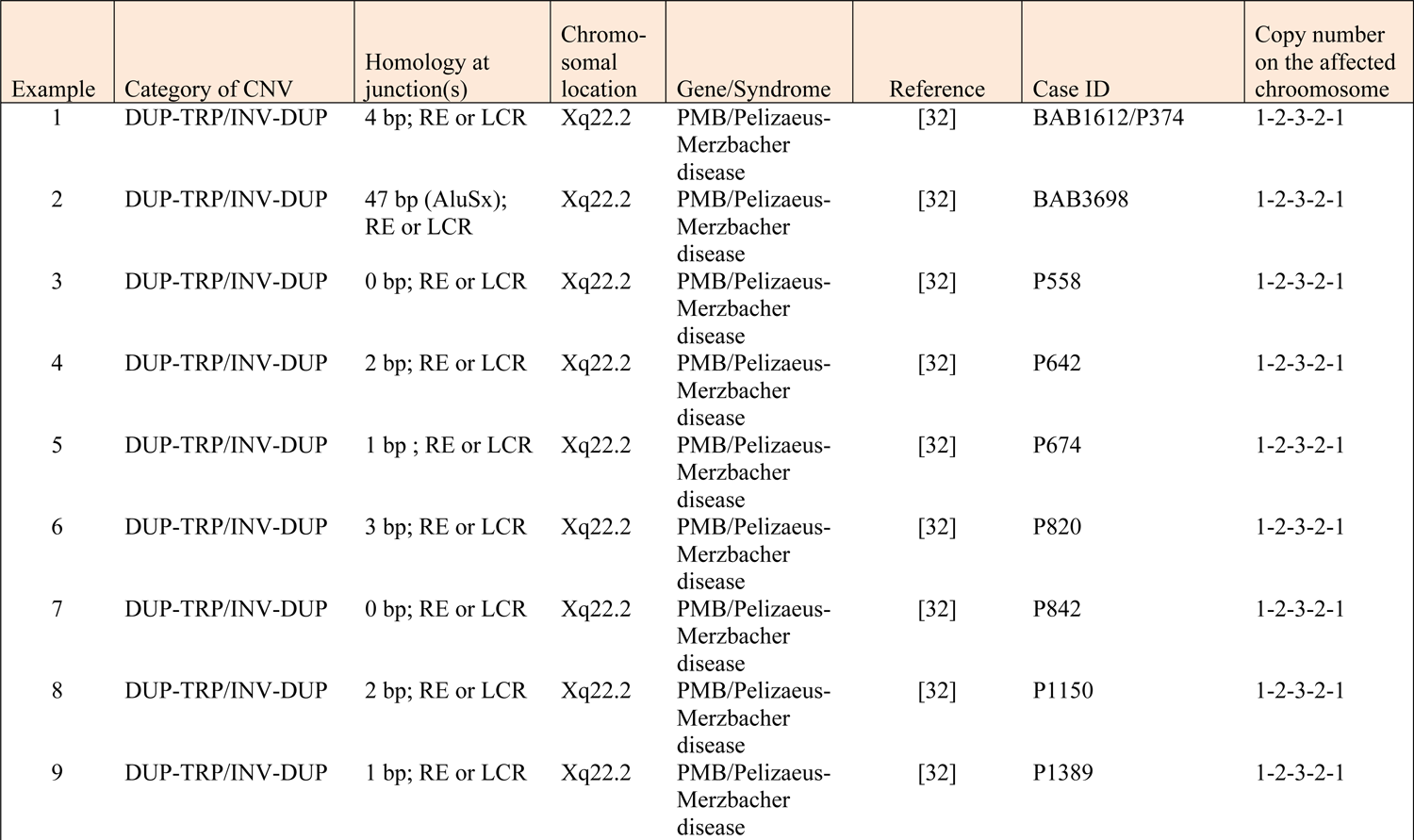
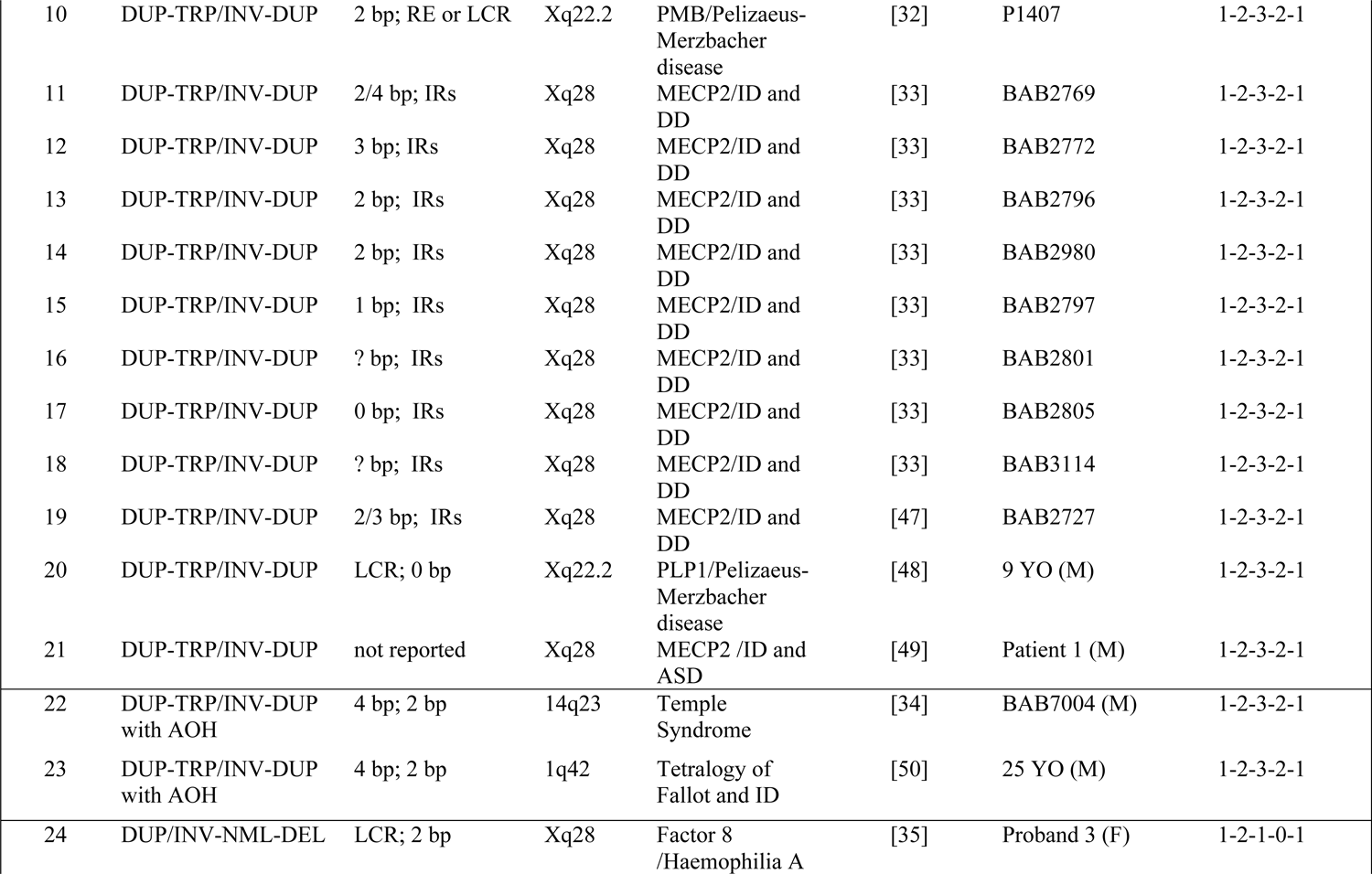
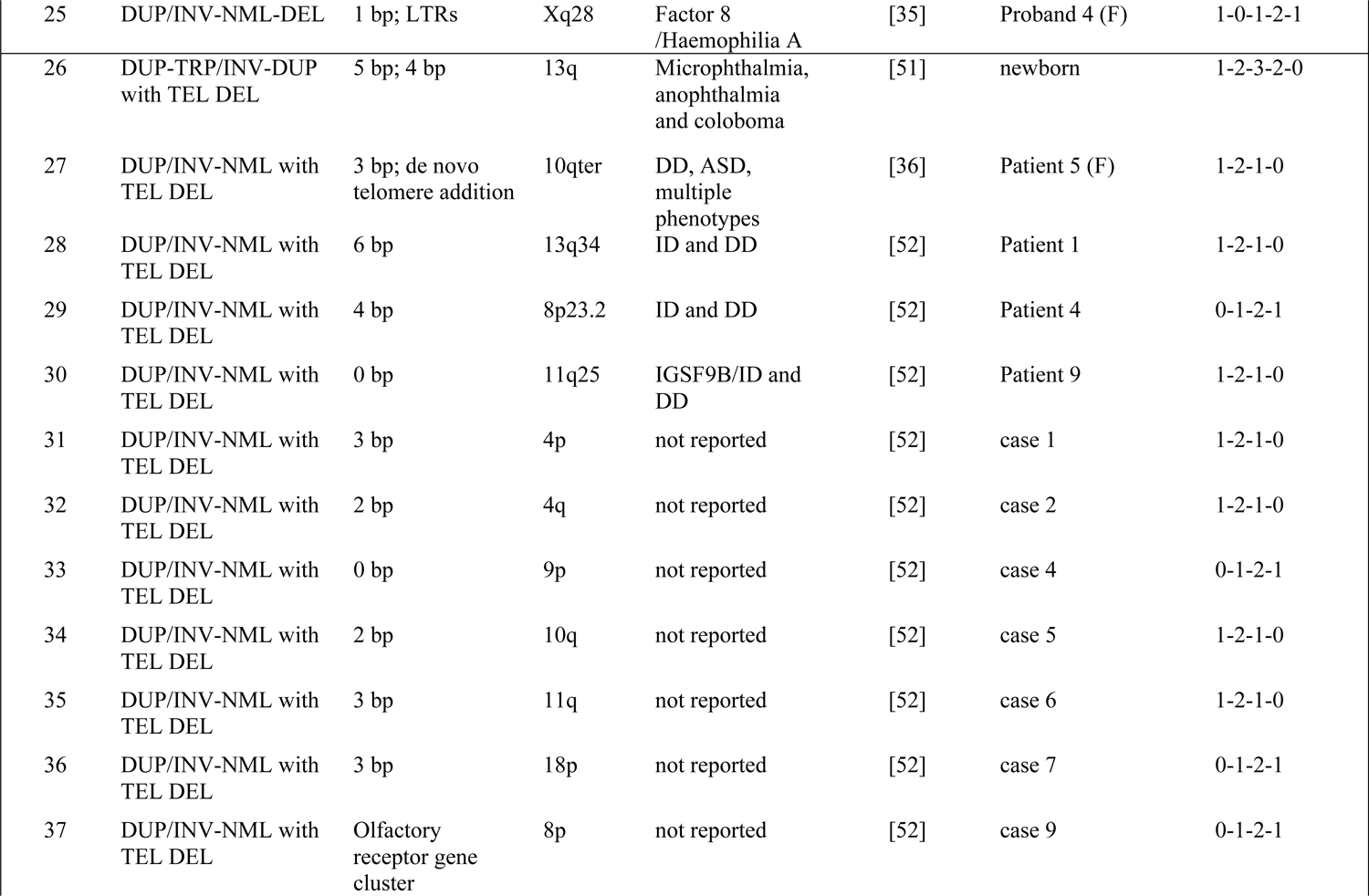
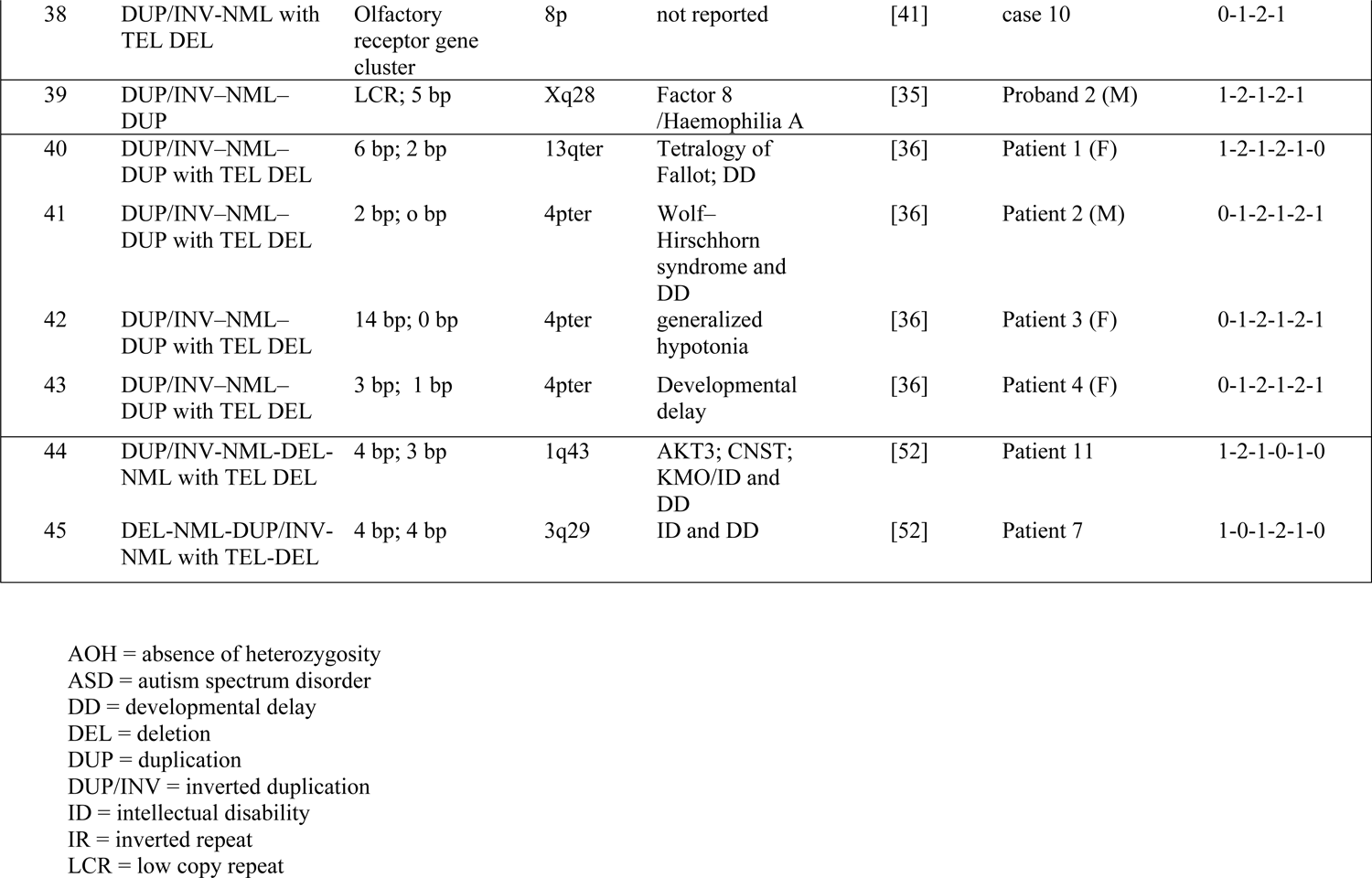
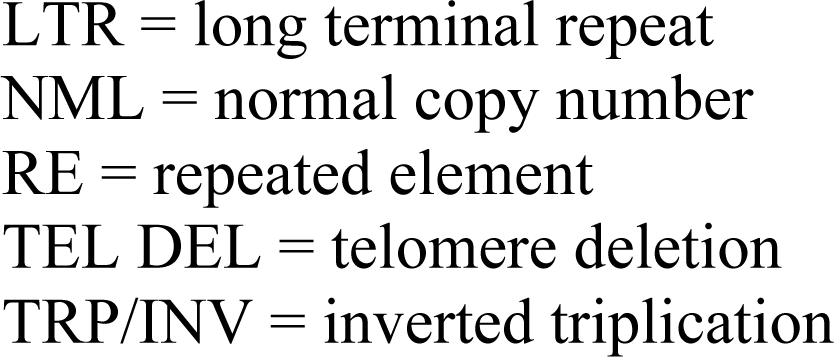
Examples from the human literature of categories of local inverted SVs.

Some DUP-TRP/INV-DUP events are flanked by a region that has become homozygous (AOH). These cases have allele compositions within and adjacent to the duplicated region, often extending through the adjacent telomere, that are consistent with a mitotic recombination event between the two homologues followed by segregation in mitosis (Figure 3E-G; Table 1 examples 22-23). In these cases authors invoke FoSTeS as the mechanism for this type of event, suggesting that the 3’ end of the lagging strand from the stalled fork visits the oppositely oriented repeat to continue synthesis before jumping to the homologue to complete replication of the chromosome (e.g., [34]), all occurring within in a single division cycle. In contrast, we are suggesting that homologue exchange could be an outcome of the reduction of one of the palindromic arms or could be an independent event that is executed at a subsequent division cycle.

### Inverted duplications associated with deletions

The dense distribution of repetitive elements in the human genome can precipitate situations where inverted junctions at a distance can lead to other types of secondary outcomes. When the repeats are more widely spaced (left-most orange arrow in Figure 4A, some distance from the pair of closely spaced inverted repeats diagrammed in Figure 3), then other potential recombination partners can be involved in the rearrangement of the palindrome. These duplications are illustrated using NAHR at repetitive elements, but similar structures could be generated at non-repetitive sequences through repair of DNA breaks by NHEJ, MMEJ or MMBIR. This particular outcome has recently been reported at the Factor 8 locus where NAHR at long, highly homologous inverted repeats generates one of the junctions and the second junction occurs at regions of little to no homology [35]. It produces the pattern of a deletion and inverted duplication separated by a stretch of copy-neutral DNA in between (1-0-1-2-1; Figure 4A; Figure 5; Table 1, examples 24-25, 39, 44-45).

**Figure 4:**
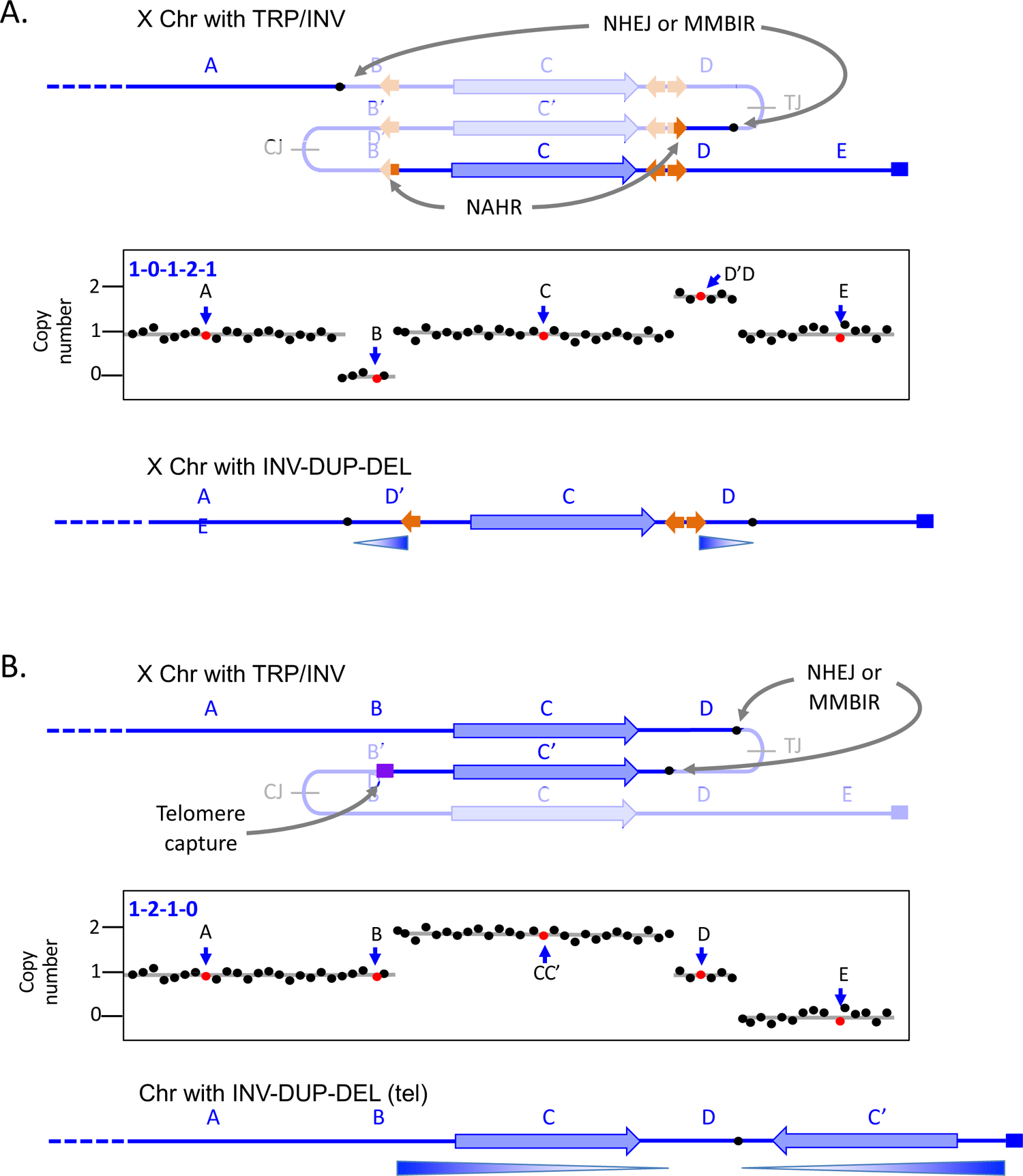
Direct and inverted duplications. A) In this example, modeled after the Factor 8 locus on the X chromosome in humans, additional repeats provide other opportunities for rearrangements of the triplicated locus that remove the centromere-proximal junction. The telomeric-proximal junction is removed by NHEJ or MMBIR at short regions of microhomology. This pattern of palindrome erosion results in a deletion and an inverted duplication separated by a copy-neutral segment of chromosome. B) A failed recombination/MMBIR attempt to erode the centromere-proximal junction leaves a dsDNA break that acquires or captures a new telomere. It results in the complete loss of sequences from the point of the inverted duplication to the end of the chromosome. See Figure S1 for alternate illustrations of the two rearrangement events.

**Figure 5:**
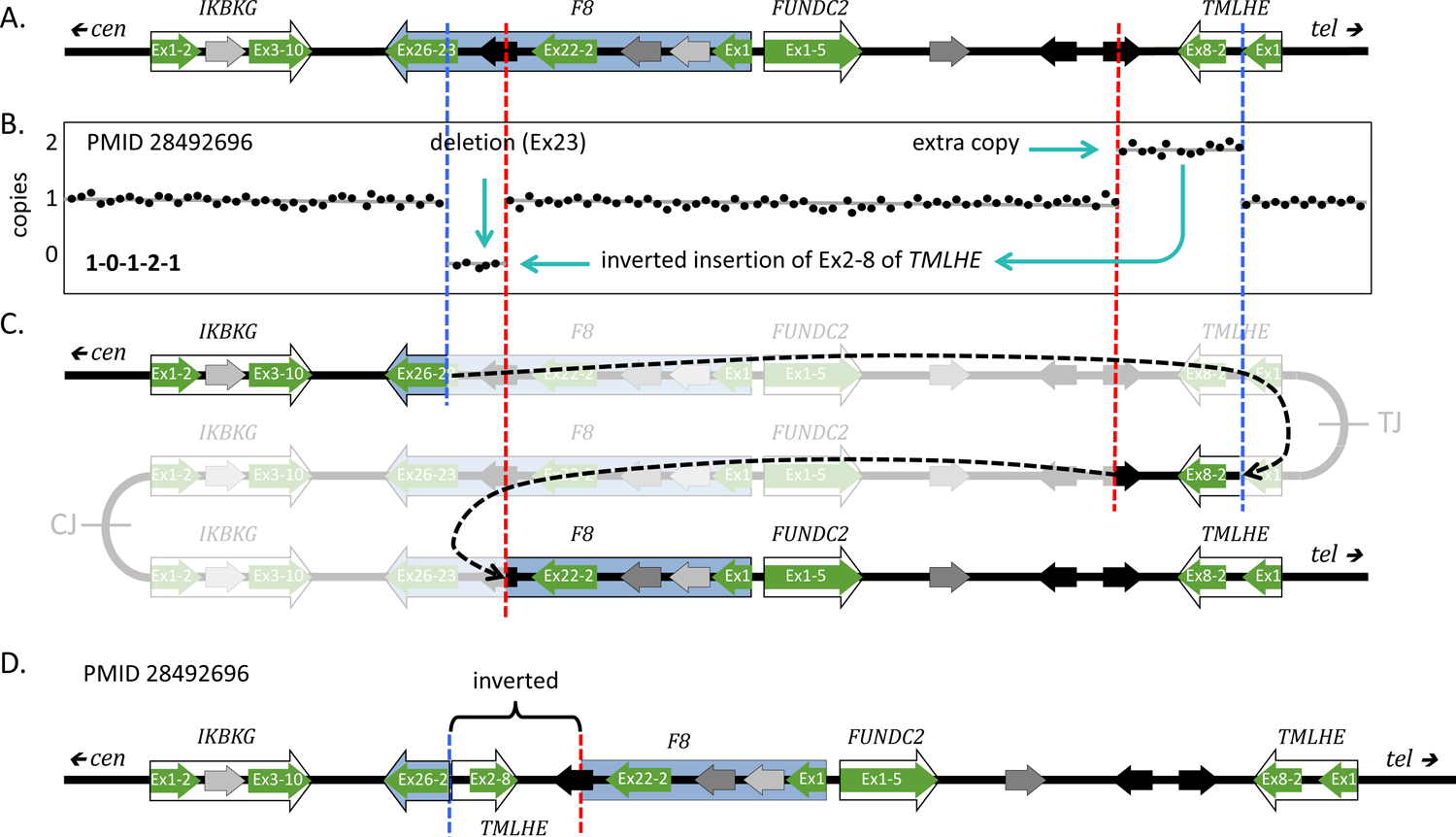
Rearrangement of an inverted triplication gives rise to a Factor 8 mutation. A) The original sequence near the Xq telomere; F8 is the Factor 8 gene, highlighted in blue. Diagram is adapted from [35]. Exons of three adjacent genes (green arrows) and three different low-copy repeats (LCRs, black and gray arrows) are highlighted. B) Stylized aCGH data for the male patient with Factor 8 deficiency (reported in [35] as PMID 28492696). Cyan arrows indicate the deleted and inverted duplicate regions. C) Hypothesized initial inverted triplication with CT and TJ palindromic junctions. Red and blue dashed lines indicate rearrangement junctions produced through NHEJ or MMEJ and NAHR at the two oppositely-oriented black LCRs. Black dashed arrows indicate the joining events; the grayed-out regions indicate the regions lost during the secondary rearrangements. D) The final structure of the F8 region of the patient in PMID 28492696 with a deletion of exon 23 and an inverted duplication of exons 2-8 of *TMLHE* and flanking regions.

The final example, also illustrated as occurring on the X chromosome, is a common form of human SV that occurs in telomeric regions. The telomere-proximal palindrome is resolved by NAHR, NHEJ or MMEJ, but the centromere-proximal palindromic junction is lost when a break is capped by a new telomere or captures a telomere from another chromosome end (Figure 4B; Table 1, examples 26-38, 40-45). Shimojima Yamamoto et al. [36] described an example of an inverted duplication on chromosome 10 that was accompanied by the loss of one of the palindromic arms, the presumptive telomere-proximal junction and all distal sequences.

### Source of the initiating TRP/INV structures

We propose that all of the above CNVs share the same starting point: a chromosome with a segment present as an inverted triplication where the center copy is inverted between two directly repeated segments with palindromes at the junctions (Figures 3 and 4). All of the different inverted CNVs can be created through rearrangements of the TRP/INV CNVs by well-characterized pathways, but what is the mechanism for forming the initial, inverted triplications and what is the nature of the initial palindrome?

We have been investigating the amplification of the yeast *SUL1* locus when cells are grown under conditions of limiting sulfate [11, 13, 37], and have acquired evidence that a replication error involving template switching of the leading strand to the lagging strand template at a replication fork is the initiating event in the amplification. Over the course of 50-200 generations, each independent culture gives rise to one or a few variants with increased fitness that contain an interstitial inverted triplication of *SUL1*, the adjacent origin of replication (*ARS228*) and centromere- and telomere-proximal palindromic junctions mapping to pre-existing short interrupted inverted repeats (Figure 6A; CJ and TJ). Analysis of nearly one hundred junction sequences [13] reveals that the repeats involved in forming the inverted triplication are quite small (between 4-8 nt) with an optimal spacing of 40-80 bp. There are very few recurrent sites but more than 95% of the junctions cluster within the *ARS228* replicon [13] and in strains deleted for *ARS228* larger amplicons are recovered that include an adjacent replication origin [11]. These results suggest that replication is involved in the formation of the inverted triplications. In a genetic screen to detect amplification of the *SUL1* region without the use of chemostats, we find that introducing nearby dsDNA breaks doesn’t increase the frequency of inverted products, suggesting that the initiating event is not a double-strand DNA break [13].

**Figure 6:**
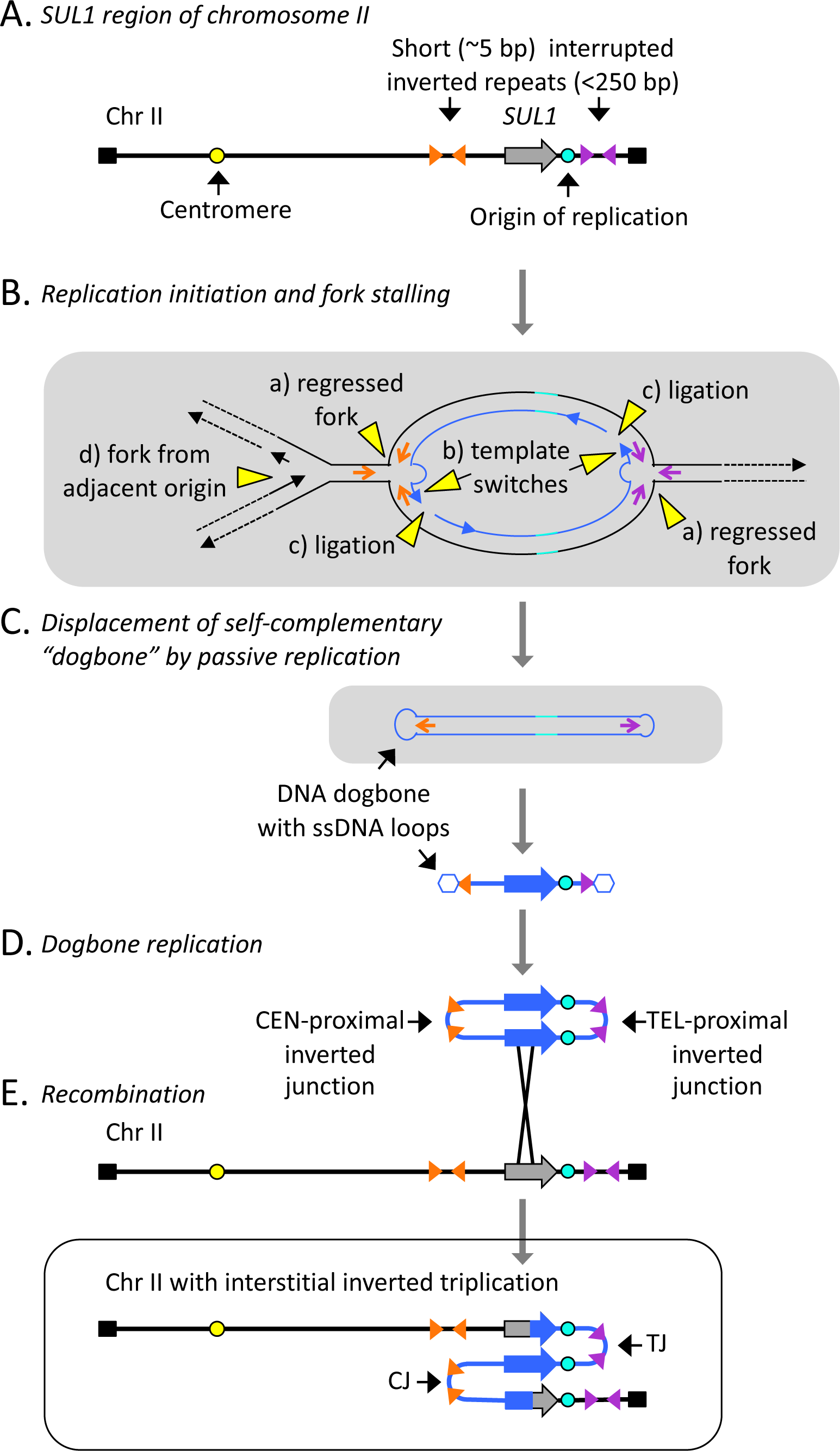
Mechanism that generates an inverted triplication of the *SUL1* region in yeast (ODIRA)—modified from Figure 1 in Martin et al, 2023 [13]. A) The structure of the *SUL1* locus in yeast includes an origin of replication (*ARS228*) and adjacent short (∼5 nt) inverted repeats separated by 40-80 nt of sequence. These interrupted inverted repeats mark the sites of the centromere- and telomere-proximal junctions (CJ and TJ, respectively) after inverted triplication. B) Process of template switching between the leading strand and its migration to complementary sequences on the lagging strand template. After extension of the 3’ end of the leading strand, it becomes ligated to the adjacent Okazaki fragment. If both forks undergo the same event, it results in a closed loop of self-complementary DNA. C) Displacement of the self-complementary loop can be achieved by branch migration ahead of an incoming replication fork. D) The resulting expelled molecule (dogbone) can replicate in the next cell cycle to produce a dimeric circular molecule with two copies of the expelled DNA in inverted orientation with junctions formed from the two inverted repeats (CJ and TJ). E) In a subsequent cell cycle, recombination of the dimeric circle into the *SUL1* region of the chromosome leads to the inverted triplication. Note that the junctions CJ and TJ retain the sequence of the two short inverted repeats where the template switching occurred.

All of our results are consistent with a replication-based model for the formation of inverted triplications that involves template switching of the leading strand onto the lagging template at a stalled replication fork (Figure 6B). Once the leading strand is ligated to the adjacent Okazaki fragment at a pair of diverging replication forks, a closed, self-complementary single-stranded intermediate is removed from the chromosome by an incoming fork and is later replicated to form an inverted dimeric circular intermediate that contains the CJ and TJ sequences (Figure 6C-E). Homologous recombination with the intact chromosomal locus produces the inverted triplication (Figure 6F). This region of the yeast genome contains no significant repeated sequences (such as Tys, deltas, or tRNA genes) so the options for eroding an arm of either palindrome are limited. However, in rare cases, we recover new junctions produced through regions of microhomology. Because the model is dependent on the presence of an origin of replication in the amplified segment and on short inverted repeats, we have named this mechanism Origin-Dependent Inverted-Repeat Amplification (ODIRA; [12]). In yeast, we have new evidence that the template switches at the two divergent replication forks can occur in different cell cycles, but after recombination with the chromosome, they produce identical inverted triplication products [13].

We propose that this same aberrant replication pathway occurs in the human genome and generates the TRP/INV substrates that give rise to all manner of local inverted CNVs (Figure 1). In Figures 3, 4 and 6, the extrachromosomal inverted duplicated circular intermediate that arose through ODIRA is shown as re-integrating into the chromosome from which it arose. However, it is also possible for the intermediate to integrate into the homologue, producing a triplication with a 2:1 ratio of SNPs from the two homologues. When this event occurs in the germ line, it is possible to see the contribution from both homologues of that parent in the CNV. While most studies do not go into this level of analysis, some reports of human inverted triplications have this 2:1 ratio of SNPs from the contributing parent (for example, [38–40]).

## Discussion

Inverted duplications that end in terminal deletions are easily and economically explained by Barbara McClintock’s breakage-fusion-bridge (BFB) model [2, 41] but are also compatible with replication-based mechanisms [42]. However, interstitial inverted SVs where the end of the chromosome remains intact require another explanation as the sister chromatid fusion that occurs during BFB results in the loss of sequences distal to the point of fusion. All inverted CNVs presented in this work (including inverted duplications with terminal deletion) can be explained by secondary rearrangements of inverted triplications produced through a strand switching mechanism between the two strands at a replication fork. It is the very nature of the palindromic structure of the inverted triplications that makes them prone to rearrangement [23]: in the course of subsequent replication cycles, processes such as NHEJ, MMEJ, NAHR, or MMBIR can repair breaks arising from the unstable palindromes.

We propose that the initial formation of inverted triplications occurs by a replication error in which the leading strand at a replication fork switches to the lagging strand template at very short, interrupted inverted repeats [12] which are found at very high density in all genomes. The size of the interruptions at ODIRA junctions in yeast is consistent with the length of the single stranded gaps between Okazaki fragments on the lagging strand [13]. The presence of a single stranded gap on the lagging strand provides the opportunity for the leading strand to switch to the lagging strand template. As Okazaki fragment size is the same across eukaryotes, we would expect to find similar junctions in human inverted triplications. All of the TRP/INV junctions we found in the literature were large enough to include multiple replication origins but had considerably larger interruptions in their palindromic arms, similar to the secondary rearrangements we find in a subset of yeast junctions. These results suggest that in the human examples, the palindromic junctions had already undergone secondary rearrangements to increase the size of the interruption to the point that the palindromic arms were no longer inducing unstable secondary structures.

The Fork Stalling and Template Switching mechanism (FoSTeS [43]) has also been suggested as a mechanism for various forms of SVs. The model was inspired by work in *E. coli* where Lac+ direct amplicons arise under conditions of stress. Mutations implicate the flap endonuclease function of DNA polymerase I and the lagging strand template [44] in their formation. The model, devised to account for the microhomology at the novel junctions, proposes that the 3’ end of an Okazaki fragment makes the jump between the *E. coli* chromosome and the F’ conjugating plasmid and back again to the chromosome. But in eukaryotes, because Okazaki fragments are only ∼150 bp (>10X times shorter than is found in *E. coli*), it is perhaps surprising that the 3’ end of such a short strand can sequentially invade a different replication fork, dissociate from it after synthesizing long stretches of DNA and then return to the original fork before completing the nascent lagging strand, all within a single S phase. Biochemically, the lagging strand template is in close proximity to the 3’end of the leading strand, making template switching within a single fork a more energetically feasible mechanism compared to FoSTeS. Despite FoSTeS being widely cited as a likely explanation for various inverted segmental variants in humans, we have not been able to find any published accounts with corroborating experimental evidence in model eukaryotes for FoSTeS. One report of intrachromosomal template switching (ICTS) in yeast by Tsaponina and Haber [45] shares some similarities with FoSTeS but differs in that the length of sequences synthesized after the jump to a new chromosomal site is very short. In addition, they did not investigate which strand at the replication fork was involved in the template switch.

In addition to providing a unifying model for a variety of congenital inverted SVs in humans we propose that ODIRA may also be a significant pathway to forming SVs in cancer cells and may provide an experimental system to dissect this mechanism. Reports of genome-wide palindrome analysis reveal a rise in palindromic sequences in a variety of cancers [19–24] but the length of spacers and their association with amplified segments remains unexplored. Are these palindromes evidence of inverted triplications? Are they a cause or consequence of some other process that is permissive for genome rearrangements? If they are formed through the same replication error we are proposing for yeast *SUL1*, then what microenvironment/stress conditions cause the aberrant template switching and are there interventions that can block these replication products or their processing? Long-read sequencing data on tumors in early stages of their development, as well as genetic dissection in models such as yeast, will be invaluable for answering these important questions.

## Acknowledgements

We are grateful for the discussions with and thoughtful comments of our colleagues Elizabeth Kwan, Rebecca Martin, Gina Alvino, Amy Moore, Joe Armstrong, Evan Eichler, and Xavi Guitart. The work was supported by National Institutes of Health grants to BJB (R01 GM018926 and R35 GM122497) and MJD (P41 GM103533) and a National Science Foundation grant to MJD (1120425). We apologize in advance for overlooking anyone’s work that is relevant to the discussion here and encourage you to contact BJB with your thoughts and comments. There were many more examples that could have been included in our analysis but we limited our analysis to those that had well characterized junctions. Again we apologize and look forward to hearing more about these other patients.

## Supplementary Figure

**Figure S1:**
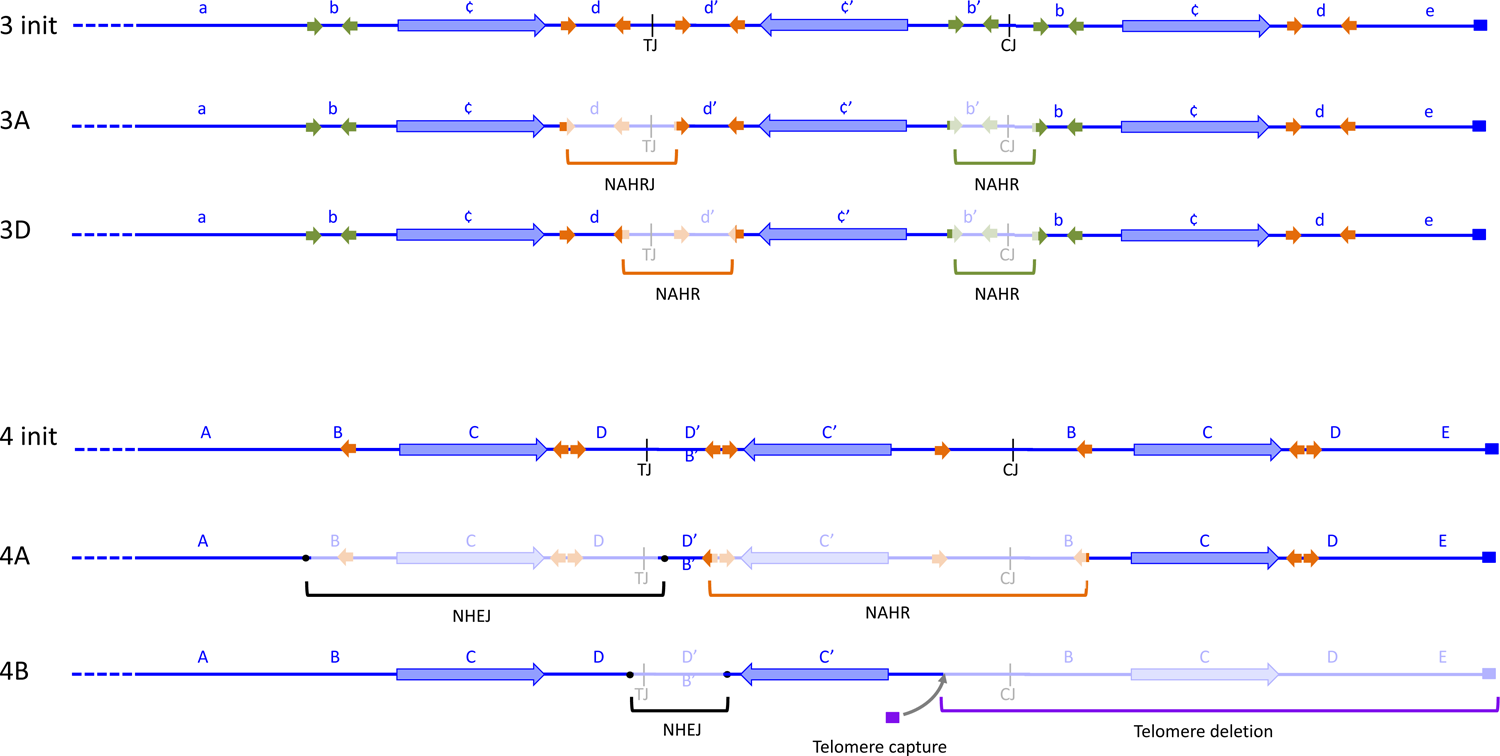
Alternate depictions for secondary rearrangements of inverted triplications. 3init, 3A and 3D refer to images from Figure 3. Init = inverted triplication before rearrangement. 4init, 4A and 4B refer to images from Figure 4. Init = inverted triplication before rearrangement. Brackets indicate rearrangement junctions created by NAHR, NHEJ or MMBIR. The grayed-out regions indicate the regions of the inverted triplication that are deleted during the secondary rearrangements.

